# A novel sperm-derived seminal fluid protein in *Caenorhabditis* nematodes

**DOI:** 10.1101/2022.09.22.509081

**Authors:** Katja R. Kasimatis, Christine Rehaluk, Locke Rowe, Asher D. Cutter

## Abstract

Nematode sperm contain subcellular vesicles known as membranous organelles (MOs) that fuse with the cell membrane upon sperm activation to release their soluble contents into the extracellular space. The second most abundant proteins in the MOs belong to the conserved Nematode-Specific Peptide family, group F (NSPF) gene family. We hypothesize that these proteins contribute to seminal fluid and are part of post-insemination reproductive tract dynamics. We characterized the anatomical region where the NSPF proteins likely function during fertilization using whole-worm immunostaining of a His-tagged *nspf-1* transgene. We confirmed that NSPF proteins are transferred to females during mating. NSPF proteins localize to the uterus lumen when transferred to mated females and in unmated adult hermaphrodites. These results suggest that the uterine localization of the NSPF proteins is likely a functional property of both male-derived sperm and self-sperm and not incidental to the point of transfer during mating. In males, we found that NSPF presence and abundance was correlated with reproductive maturity. We then used experimental evolution to compete the wildtype allele against a deletion allele in 10 replicate obligate-outcrossing populations. We calculated a mean selective disadvantage of 0.1% for the deletion allele, which indicated that the NSPF genes are beneficial to male fitness. This conclusion was reinforced by qualitative trends from lower powered single-generation fertility assays. Together we demonstrate that nematodes use a novel mechanism for generating seminal fluid proteins and show that the highly abundant NSPF proteins likely have a beneficial impact on fitness.

## INTRODUCTION

Post-insemination reproduction studies have largely focused on interactions mediated by seminal fluid. Seminal fluid is a complex mixture of cells, proteins, and other small molecules that are transferred to the female along with sperm (Poiani, 2006; Perry *et al*., 2013). As a whole, seminal fluid is critical for male fertility – sperm alone is not enough to ensure fertilization – and is implicated in complex behaviors, such as sperm competition, male-female signaling and sexual conflict (Poiani, 2006; McDonough *et al*., 2016). The study of seminal fluid proteins by evolutionary biologists has largely taken a narrow focus on the study of dozens of *Drosophila* accessory gland proteins (Acps) (Chapman, 2011). The genes underlying many of these Acps are characterized by their elevated evolutionary rate relative to non-reproductive proteins, indicating adaptive evolution through sexually antagonistic coevolution, but also sperm competition (Begun *et al*., 2000; Swanson & Vacquier, 2002; Clark *et al*., 2006; Wilburn & Swanson, 2016) or relaxed selection (Dapper & Wade, 2016). However, the null expectation for the evolutionary histories of seminal fluid proteins is not well-developed. Employing new approaches and new taxa may provide a more precise, complete, and generalizable understanding of the complex roles, evolutionary histories, and fitness effects of seminal fluid proteins (Rowe *et al*., 2018; Hopkins & Perry, 2022).

*Caenorhabditis* nematodes are a powerful model system for discovering new roles for seminal fluid proteins and expanding our view on the null expectations for their evolutionary histories. First, mating system differs across the genus, with *C. elegans* representing a lineage transition from obligate outcrossing to hermaphroditic self-fertilization (Baldi *et al*., 2009). In *C. elegans*, males occur in approximately 0.1% of selfed progeny due to X chromosome non-disjunction events, and natural populations show evidence of some outcrossing (Andersen *et al*., 2012; Félix & Duveau, 2012; Richaud *et al*., 2018). Both the sperm that derives from males and hermaphrodites are stored in the same region of the gonad of hermaphrodites, allowing for direct competition of sperm types (LaMunyon & Ward, 1998). Using the genetic tools developed in *C. elegans*, researchers can manipulate hermaphrodites to develop as pseudo-females that rely on males to reproduce, which allows direct comparison of the genetics and fitness effects between self-sperm and male-derived sperm to disentangle conflict and cooperation. Additionally, broader contrasts can be made among closely related species with different mating systems and therefore varying opportunities for sexual conflict (Ting *et al*., 2014; Yin *et al*., 2018). Second, the small size and rapid generation time of nematodes make them amenable to fitness assays within and across multiple generations. Multi-generation assays allow for detection and quantification of even small fitness effects. Finally, nematodes have unique amoeboid-like crawling sperm which can be visualized through the transparent cuticle, allowing for spectacular live visualization of sperm and seminal fluid dynamics inside of females or hermaphrodites (Smith & Stanfield, 2011; Ting *et al*., 2014; Hansen *et al*., 2015; Yin *et al*., 2018). Very few seminal fluid proteins, however, have been identified and functionally examined in *Caenorhabditis* (Smith & Stanfield, 2011; Yin *et al*., 2018). Moreover, unlike males in most insect and mammalian systems, nematode males do not have accessory glands specific to seminal fluid production (McGraw *et al*., 2015), and the anatomical region in which seminal fluid proteins are produced is still unknown. Thus, this system provides an underexploited opportunity for discovering new features of reproductive evolution that can be evaluated and, perhaps, challenge existing paradigms.

An intriguing component of nematode sperm cells are subcellular vesicles known as membranous organelles (MOs). In inactive spermatids, MOs are located around the periphery of the cell (Ellis & Stanfield, 2014). Upon ejaculation, spermatids are released from the seminal vesicle and undergo activation as they pass through the vas deferens. During activation, MOs fuse with the cell membrane and create cup-like structures as well as releasing their soluble protein contents into the extracellular space (Ellis & Stanfield, 2014). No accessory gland-like tissues have been identified in *Caenorhabditis* and the source – as well as contents – of seminal fluid are largely unknown (except see Smith & Stanfield, 2011). Consequently, we hypothesize that these MO fusions provide a novel contribution to the seminal fluid proteome. Characterization of the MO proteome identified the Nematode-Specific Peptide family, group F (NSPF) as the second most abundant set of proteins in the MO (Kasimatis *et al*., 2018). Kasimatis *et al*. (2018) additionally showed that the NSPF genes are conserved in both sequence identity and synteny across the genus. Transcriptomic data support the molecular evolution analyses by capturing NSPF expression in *C. elegans* males along with NSPF ortholog expression in *C. remanei* males (Albritton *et al*., 2014). High-resolution spatial gene expression mapping localizes NSPF expression to the proximal gonad of adult *C. elegans* males (Ebbing *et al*., 2018). Interestingly, transcriptomic data also identify expression in *C. elegans* hermaphrodites but not *C. briggsae* hermaphrodites, suggesting that this gene family has been incorporated into in at least some self-sperm pathways (Albritton *et al*., 2014).

Here we directly test the hypothesis that MO fusion represents a novel mechanism for contributing proteins to seminal fluid. We use immunostaining of the abundant NSPF proteins as a marker to identify the functional regions that NSPF proteins are associated within males and females/hermaphrodites. We then investigate the impact on fitness of the NSPF gene family using a combination of within generation fertility assays and 20 generations of experimental evolution. Our results indicate that the NSPF proteins are a component of seminal fluid with a marginally-detectable male fitness benefit and imply that even a highly abundant, conserved seminal fluid protein can exert only a subtle function.

## MATERIALS AND METHODS

### Molecular biology

To create a tagged *nspf-1* gene, guides targeting the sequence in the same intergenic region utilized by the ttTi5650 MosSCI site have previously been described (Dickinson *et al*., 2015) and inserted into pDD162 (Addgene #47549) to make plasmid pMS4. The repair template plasmid (pKK4) was assembled using the NEBuild HiFi Kit (NEB) from a combination of a restriction digest fragment and a synthetic gBlock Gene Fragment (IDT). The gBlock included the 647 base pairs upstream of *nspf-1* as the promoter region, the *nspf-1* coding sequence with six histidines added before the stop codon, and the *tbb-2* 3’ UTR. The plasmid was purified using the GeneJET Plasmid Miniprep Kit (Thermo Scientific) and assembly junctions were verified by Sanger sequencing. A complete list of primers can be found in Table S1.

### Strain generation

All strains used in this study are listed in Table S2. The NSPF genes were previously deleted using CRISPR/Cas9 to create strain PX814 (fxDf1 (II: 2,484,339 - 2,487,244)) (Kasimatis *et al*., 2018). To generate an obligate outcrossing deletion strain, JK574 (*fog-2*(q71) V) was crossed with PX814 and then backcrossed 5 times with JK574 to create the obligate outcrossing, deletion strain VX194. The deletion region was verified by Sanger sequencing.

Insertion of the His-tagged *nspf-1* transgene was done by CRISPR/Cas9 using standard methods (Dickinson *et al*., 2015). Briefly, a mixture of 10ng/μL repair template plasmid, 50ng/μL plasmid encoding CAS9 and the guide RNA, and 2.5ng/μL of the co-injection plasmid pZCS16 was injected into the gonad of young adult hermaphrodites from PX814. Hygromycin B (BioShop) was added to the injection plates at a final concentration of 250μg/mL three days after injection and hygromycin resistance (HygR) was used as a selectable event. Worms were kept at 25°C for the first five days immediately following injections and then transferred for 20°C for subsequent generations. Successful insertion was confirmed by PCR (Table S1) and Sanger sequencing. Worms were then backcrossed five times to PX814 to create the final strain VX300 (yf*Si*1[*nspf-1*p::*nspf-1*::6xHis::*tbb-2* 3’ UTR + loxP, II:8420157]). We tagged the C-terminal of *nspf-1* since these proteins are predicted signaling peptides (Kasimatis *et al*., 2018) and, thus, the C-terminal is likely included in the mature peptide. The mature peptides for *nspf-1* and *nspf-2* differ by two amino acids and differ by four amino acids between *nsfp-1* and *nspf-3*. These differences are not predicted to change the cleavage site or secondary structure of the peptide. Across the genus, *nspf-1* is more conserved than *nspf-3* (Kasimatis *et al*., 2018). Thus, expression of *nspf-1* likely is representative of the localization and function of this gene family.

### Whole-worm immunohistochemistry

Worms were formaldehyde-fixed and stained following a modified version of the Ruvkun and Finney protocol (WormAtlas). Briefly, worms were cultured at 20°C on 10cm NGM-agar plates containing 100ng/mL carbenicillin and seeded with formaldehyde killed *E. coli* OP50 (following Beydoun *et al*., 2021), which was used to reduce non-specific binding of the His-antibody to *E. coli* produced histidines. At each developmental stage used, populations of at least 1,000 worms were fixed in a 1% formaldehyde solution and flash frozen in dry ice with ethanol. Incorrect antibody binding was blocked using solution of: 0.3M glycine, 10mM imidazole, 1% BSA, 0.5% skim milk, and 0.25% Triton-X-100 in PBS. Worms were stained for 1 hour at room temperature using a 1/4000 dilution of Recombinant Alex Fluor 488 Anti-6x His Tag antibody (Abcam #ab237336) and co-stained with DAPI. The complete method along with buffer recipes can be found in File S1.

To examine the localization of *nspf-1* in males and hermaphrodites across development, VX300 male-rich populations (∼40% males) were age synchronized by bleaching (Kenyon, 1988) and then formaldehyde-fixed during: L3 (28 hours post-L1), L4 (40 hours post-L1), day 1 of adulthood (46 hours post-L1), day 2 of adulthood (70 hours post-L1), and day 8 of adulthood (190 hours post-L1). As a control, JK574 males and pseudo-females were concurrently cultured and fixed at the same developmental stages. In our hands, worms developed more slowly and less synchronously on plates with dead OP50 than on live OP50. Thus, the exact time in development may be asynchronous between worms, but the developmental stages were accurate. At least 100 stained worms from each strain × sex × developmental stage were scored for the presence and location of fluorescence (File S2). The localization of fluorescence was characterized as occurring in one or more of the following regions: head, distal gonad, proximal gonad (including the spermathecae or seminal vesicle), uterus/vas deferens, vulva/cloaca, or tail. Worms were categorized as “not stained” if no fluorescence was observed.

To determine whether or not males transfer NSPF-1 to females during mating, we crossed JK574 pseudo-females with VX300 males. Both strains were age-synchronized by bleaching followed by isolation of virgin pseudo-females (n = 400) and males (n = 150) during late L4 to 10cm NGM-agar plates containing 100ng/mL carbenicillin and seeded with formaldehyde-killed *E. coli* OP50 (n = 3 replicate plates). Worms were mated overnight and then formaldehyde-fixed and stained. One hundred pseudo-females were scored for the presence and location of fluorescence.

Count data for the presence of fluorescence were analyzed using a χ^2^ test based on a contingency table of strain × presence of fluorescence for each sex and developmental stage separately. Specifically, we tested the null hypothesis of no difference between counts of stained individuals and counts of unstained individuals between the control (JK574) and His-tagged (VX300) strains. All analyses were conducted using the R statistical language v4.0.0 (R Core Team, 2020).

### Dissected gonad immunohistochemistry

Testes were methanol-fixed and stained following a modified version of the Shakes *et al*. (2009) protocol. Briefly, worms were cultured at 20°C on 10cm NGM-agar plates and seeded with *E. coli* OP50. Young adult males were decapitated using a 30 gauge needle to release the gonad and then transferred to ColorFrost Plus slides (Fisher Scientific). Gonads were fixed in methanol followed by acetone. Incorrect antibody binding was blocked using solution of: 0.3M glycine, 10mM imidazole, 1% BSA, and 0.25% Triton-X-100 in PBS. Worms were stained for 1 hour at room temperature using a 1/4000 dilution of Recombinant Alex Fluor 488 Anti-6x His Tag antibody (Abcam #ab237336) and co-stained with DAPI. The complete method along with buffer recipes can be found in File S3. Image stacks were processed using ImageJ.

### Fertility Assays

We assayed the overall reproductive success of VX194 deletion males late in adulthood. We isolated virgin deletion males on day 1 of adulthood and maintained as virgins until the start of the assay. On day 5 of adulthood, individual males were mated with three wildtype, virgin females (strain JK574) for 24 hours. As a control, wildtype males (strain JK574) were mated to wildtype females following the same timing and male to female ratio. Matings were conducted on small NGM-agar plates (35 mm diameter) seeded with 10μL OP50 *Escherichia coli*. After 24 hours, each male was removed and the females were transferred to a new small plate to continue laying eggs. Females were transferred to new plates every 24 hours until progeny production ceased. The total number of progeny was counted as a measure of each male’s reproductive success (File S3). Fertility data were analyzed using R (R Core Team, 2020), with the significance of non-competitive reproductive success evaluated using Welch’s Two Sample t-test and an analysis of the power of the comparison computed. Previous fecundity data were replotted using the data available in Kasimatis et al. (2018) Additional File 9.

### Experimental evolution design and worm culture

To create a base population, reciprocal crosses of virgin females and males from strains JK574 and VX194 were performed. The F1 heterozygous progeny were divided into 10 replicate populations of 500 worms with equal numbers of each sex to start the first generation. After generation one, populations were expanded to a census size of N = 1,000 worms per replicate and evolved for 20 generations. To start each generation, age synchronized larval stage 1 (L1) worms were plated onto four 10 cm NGM-agar plates seeded with OP50 *E. coli* at 20°C with a density of 250 worms per plate (Brenner, 1974; Kenyon, 1988). Worms were left to mate for the first five days of adulthood before eggs were once again collected on day six to synchronize for the next generation. During this mating time, worms within each replicate were pooled and redistributed to fresh NGM-agar plates twice to ensure they never starved. The mating duration provided ample opportunities for females to mate with multiple males. A subset of worms from all replicates was frozen every five generations for future experiments. At generation eight all worms were frozen and the experiment paused for five months due to COVID-19 lockdown restrictions. The thawed population sizes were in excess of the census size and plated at a density of 250 worms per plate, following the regular experimental protocol.

### Amplicon sequencing and read calling

We performed amplicon sequencing on pooled samples of 500-800 L1 worms from experimental evolution generations 11 and 20. Two control samples were generated, each with approximately 700 worms: 1) F1 heterozygous progeny generated by crossing strains JK574 and VX194, and 2) pooled worms generated by mixing an equal number of JK574 and VX194 worms and sampling from this pool. The wildtype allele was amplified using forward primer (5’ to 3’) TGCAATGGATTTCTGGTGATAAC and reverse primer (5’ to 3’) TCTGGCAGTAGGAAAGTGAG, yielding a product of size 230 base pairs. The deletion allele was amplified using forward primer (5’ to 3’) AGGATAAGAGAGACCCAGAC and reverse primer (5’ to 3’) TCTGGCAGTAGGAAAGTGAG, yielding a product of size 275 base pairs. PCR products were generated using the OneTaq Quick-Load 2x PCR Master Mix with Standard Buffer (NEB) in accordance with manufacturer instructions with *C. elegans* genomic DNA. PCR products were purified using QIAquick PCR Purification kit (Qiagen) and normalized to 20ng/uL. Products were sequenced using GENEWIZ Amplicon-EZ services.

Reads for each sample were filtered using bioawk (https://github.com/lh3/bioawk) to exclude reads <50 base pairs in length and with a mean quality score <30. Reads in each sample >230 base pairs in length were then blasted to the deletion reference sequence and retained if the blast score was significant. All remaining reads were then blasted to the unique reference sequence for both the wildtype (bases 1-102) and deletion (bases 1-147) alleles. A read was called as a wildtype or deletion allele based on the best E-value and bit score (File S4). Final counts ranged from 89,721 to 132,685 reads per sample.

### Estimating selection

Read data were analyzed using R (R Core Team, 2020) and the *lme4* v.1.13 package (Bates et al. 2015). We fit five different linear models. To account for biases during PCR amplification due to different wildtype and deletion product sizes, as suggested by the mean frequency of each allele in the two control samples (*i.e.,* 72.6% wildtype and 27.4% deletion). Models 1-4 only analyzed amplicon sequence data from generations 11 and 20. Model 5 assumed the frequency of each allele to be 50% (*i.e.,* 500 copies of the wildtype allele and 500 copies of the deletion allele for a heterozygous population of N = 500), because all worms were heterozygous at the start of the experiment. Model 6 used the frequency of the deletion allele estimated from the control samples as an alternative generation 1 estimate that includes PCR bias. Model 1 fit read counts over time using a generalized linear mixed model with a binomial logistic distribution and replicate as a random effect: glmer((WT_count_, DEL_count_) ∼ generation + 1|replicate). We present the results of Models 1 and 6 below as they represent the most conservative estimate of selection. The results of all six models are represented in Table S3 and are qualitatively similar in suggesting a decrease in the frequency of the deletion allele over time.

## RESULTS

### NSPF proteins localize to the spicules in adult males

We characterized the localization of NSPF proteins during fertilization using whole-worm immunostaining of a His-tagged *nspf-1* transgene re-inserted into the deletion allele background (PX814). We scored the presence of the fluorescent antibody bound to NSPF-1::6xHis in six anatomical regions: head, distal gonad, proximal gonad, vas deferens, cloaca, and tail (Fig. 1A). Throughout male development, we found that NSPF presence and abundance was correlated with male reproductive maturity (Fig. 1B; Fig S1). During larval stage 3 (L3), the gonad is still developing and no staining was present, which supports our previous finding that NSPF production is linked with sperm production (File S2). Sperm production begins during the L4 stage and correspondingly some L4 males had staining in both the distal (n = 17 of 126) and proximal gonad (n = 1 of 126; Fig. 2A). Interestingly, 43% of L4 males had staining in the tail region (n = 55 of 126) specifically overlapping with the developing spicules, structures that facilitate mating and sperm transfer in adult males (Emmons, 2005). These patterns are significantly different from control males (χ^2^ = 175.5, df = 3, p < 0.0001), of which only one individual showed non-specific staining along the cuticle in the proximal gonad region (Fig. 2B).

**Figure 1.**
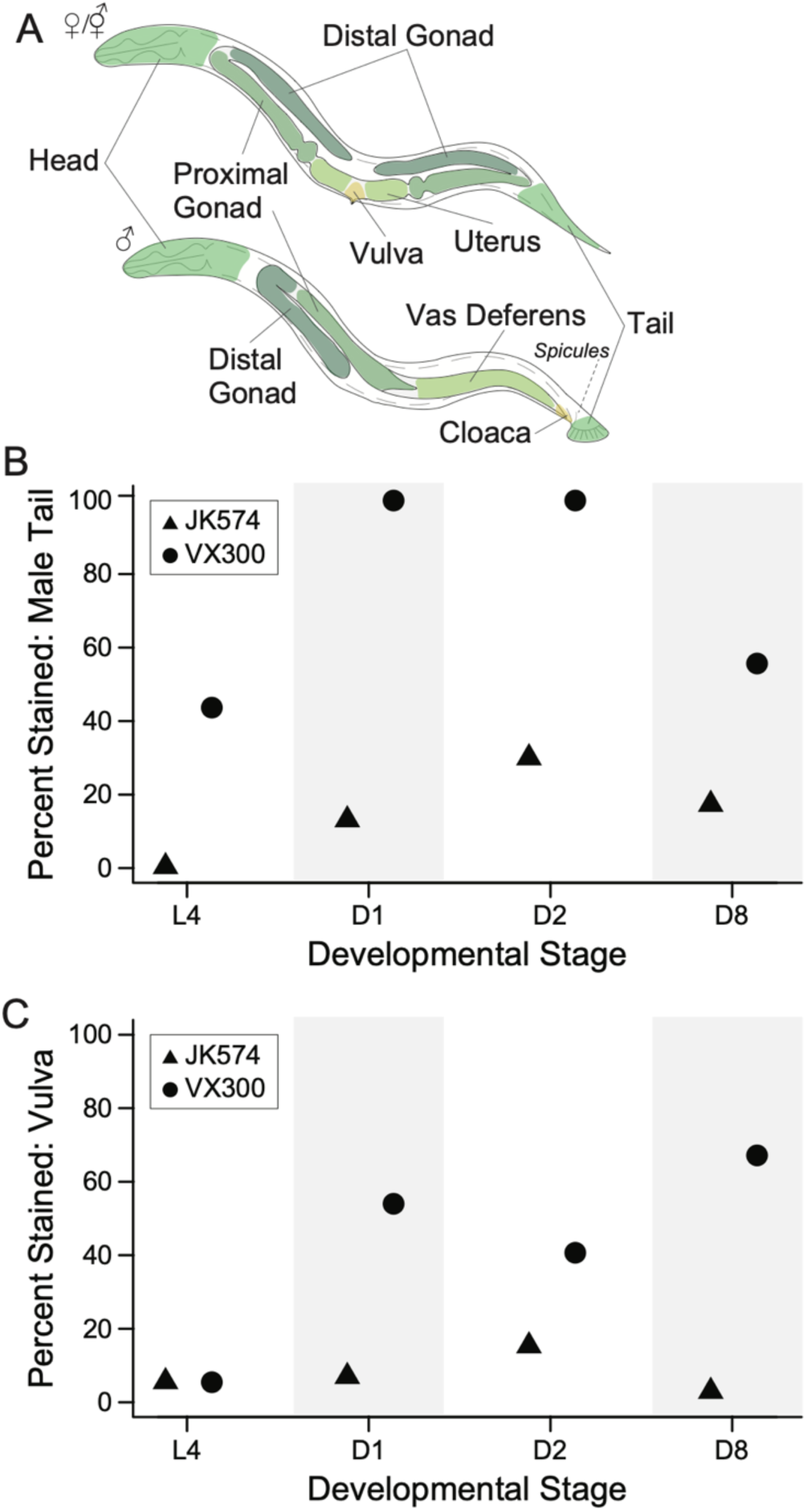
Quantification of antibody staining of 6x His-tagged NSPF-1. **A)** Diagram of the anatomical regions scored for presence of fluorescence in hermaphrodites/females and males. **B)** The percent of control males (JK574, triangle) and His-tagged males (VX300, circle) with fluorescence in the tail region during larval stage 4 (L4), day 1 of adulthood (D1), day 2 of adulthood (D2), and day 8 of adulthood (D8). **C)** The percent of control females (JK574, triangle) and His-tagged hermaphrodites (VX300, circle) with fluorescence in the vulva region during larval stage 4 (L4), day 1 of adulthood (D1), day 2 of adulthood (D2), and day 8 of adulthood (D8). Immunostaining counts across all anatomical regions is shown in Fig. S1.

**Figure 2.**
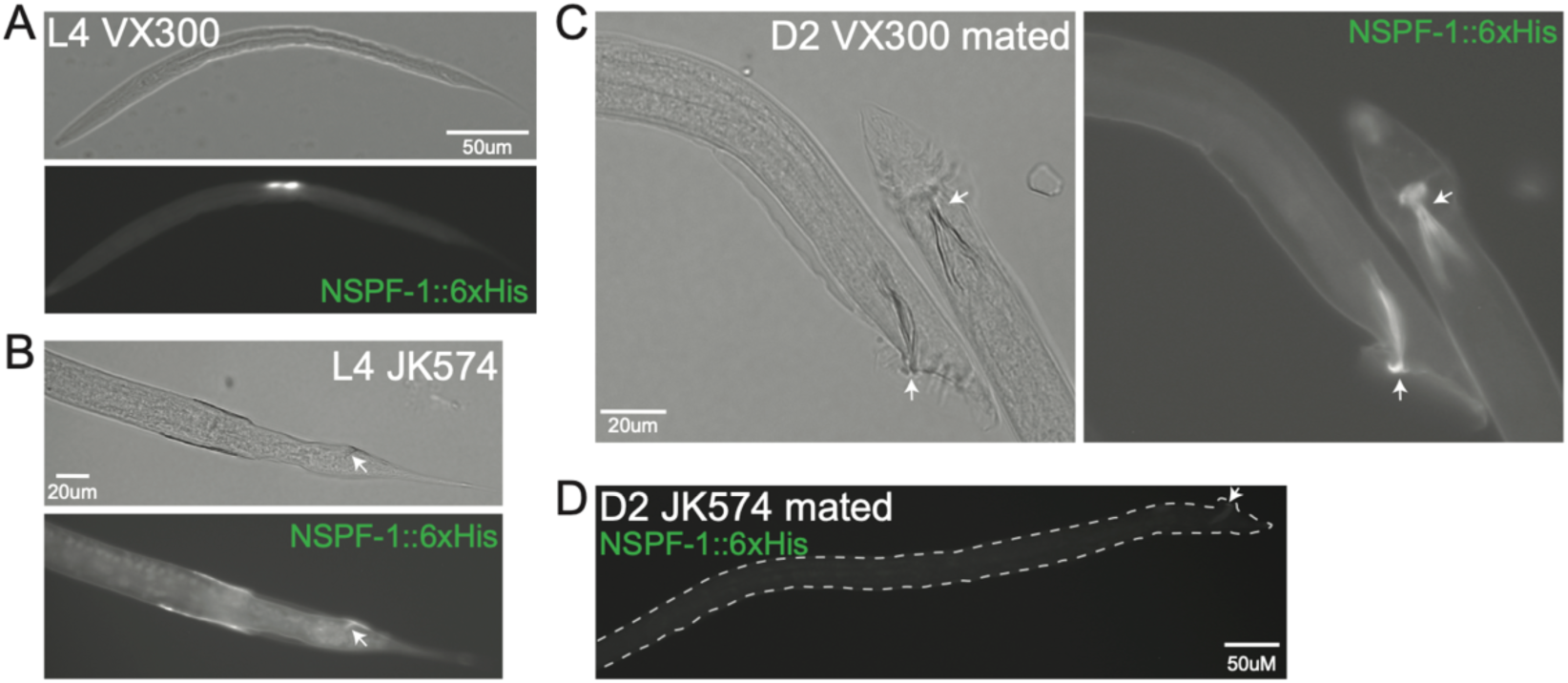
NSPF-1 localizes the spicules of adult males. **A)** Protein localized to the gonad of L4 His-tagged males. **B)** L4 control males showed off-target staining, predominantly in the cuticle including the cuticle adjacent to the spicules, but not in the spicules. **C)** In adult His-tagged males, protein localized to the spicule structure in the male tail region. **D)** In adult control males, low levels of autofluorescence are detected in the spicules. However, the signal is markedly different than seen in His-tagged males. In all images, the arrow points to the spicule.

In whole-worm staining of adult males, NSPF proteins localized solely to the spicules (Fig. 2C). Abundance appeared highest on days 1 and 2 of adulthood with 100% of males (n = 100 and 150, respectively) showing staining in the tail region (Fig. 1B). Again, these patterns were significantly different from the off-target staining or spicule auto-fluorescence observed in control males during day 1 (n = 13 of 99 stained; χ^2^ = 149.5, df = 1, p < 0.0001) and day 2 (n = 45 of 150 stained; χ^2^ = 158.5, df = 1, p < 0.0001) of adulthood (Fig. 2D). Older males, however, had a decreased signal of NSPF proteins (n = 59 of 106), though still significantly different from control males (χ^2^ = 39.4, df = 1, p < 0.0001). This pattern of protein abundance throughout development suggests that NSPF production is tightly correlated with spermatogenesis and peak reproductive output and mating activity during early adulthood.

While immunostaining of whole adult worms did not identify NSPF protein within the gonad, dissected testes of adult males confirmed localization of NSPF within spermatids (Fig. S2). The punctate patterning around the nucleus overlaps with the location of MOs, which supports the proteomic characterization of this gene family. Thus, whole worm immunostaining likely lacked the sensitivity to visualize staining within small sperm subcellular structures.

### Males transfer NSPF proteins to the female uterus during mating

To test our hypothesis that NSPF proteins form a constituent of seminal fluid, we examined whether males transferred NSPF proteins during mating. Specifically, control pseudo-females (JK574) were mated with His-tagged males (VX300) for 24 hours. Of the pseudo-females without eggs in utero, 38% (n = 8 of 21) had a clear signal of staining around the vulva and in the uterine space (Fig. 3A). This signal was significantly different from that of control females mated to control males (χ^2^ = 4.9, df = 1, p = 0.03; Fig. 3B), though given the small sample sizes false positives are possible. Only 12% (n = 15 of 129) of pseudo-females with eggs in utero showed a clear signal of staining, which was not significantly different from pseudo-females mated to control males (χ^2^ = 0.53, df = 1, p = 0.47). However, in these females the NSPF proteins appear to be in the uterus lumen, which suggests that eggs may mask the fluorescent signal (Fig. 3C). These results indicate that males transfer NSPF proteins during ejaculation and support the hypothesis that MOs contribute proteins to seminal fluid.

**Figure 3.**
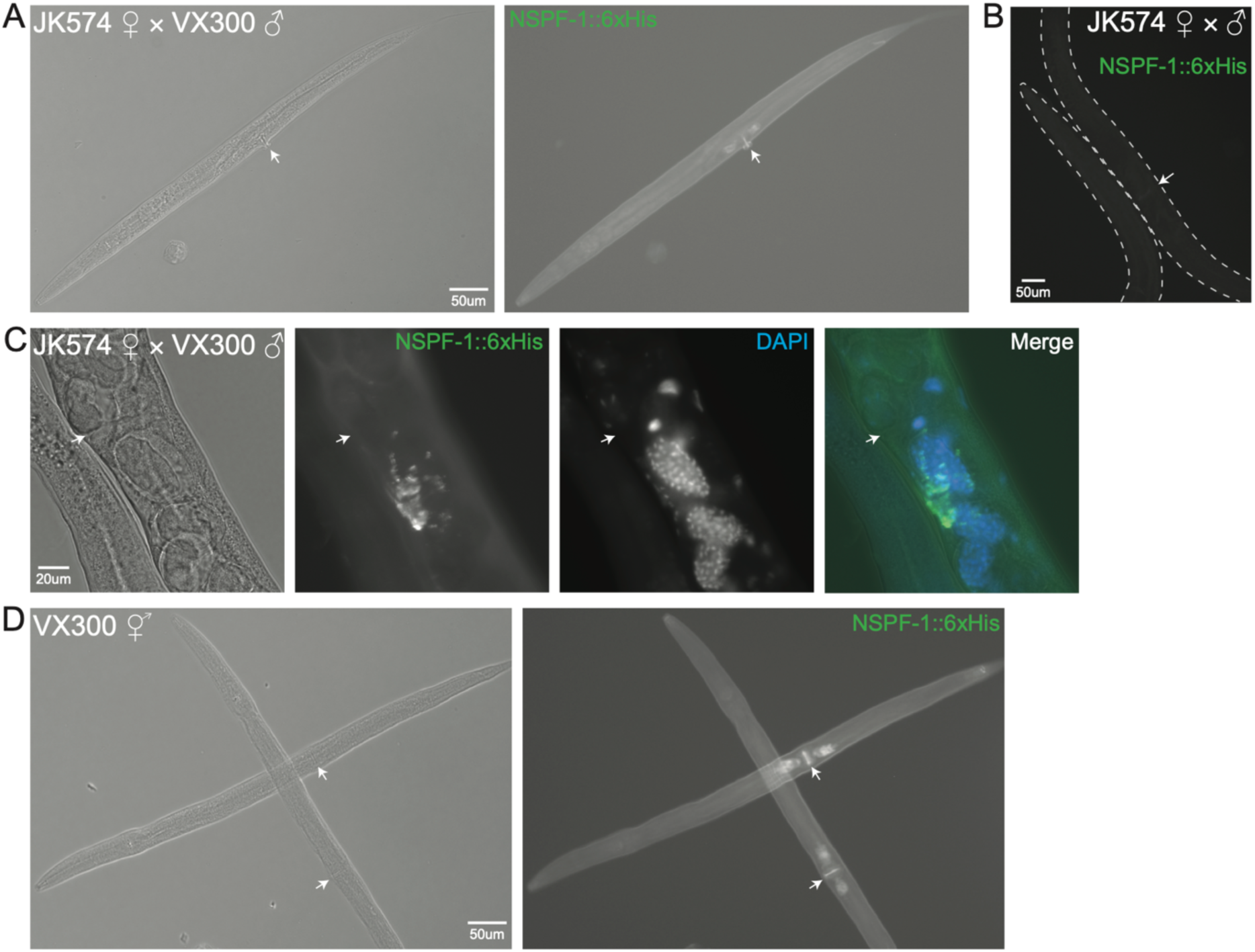
NSPF-1 localizes to the vulva and uterus of mated females and adult hermaphrodites. **A)** Control pseudo-females mated to His-tagged males showed evidence of protein localization in the vulva and uterus, especially when eggs were not yet present in the uterus. **B)** Control pseudo-females mated to control males showed no evidence of staining. **C)** Control pseudo-females mated with eggs present in the uterus after mating with His-tagged males also had protein present in the uterus, though the signal was not always clear. However, when present the proteins seemed to localize to the lumen and potentially along the surface of vulva muscle cells, as shown in the merged image. **C)** His-tagged adult hermaphrodites who had never been mated by a male showed the same protein localization in the uterus. In all images, the arrow points to the vulva.

### NSPF proteins localize to the uterus of unmated hermaphrodites

To determine if self-sperm NSPF patterns differed from male-derived sperm patterns, we characterized NSPF localization in His-tagged hermaphrodites (VX300) that had been raised in the presence of males throughout development. Again, L3 individuals showed no signal of staining (File S2). L4 hermaphrodites, however, had little to no staining and did not significantly differ from control females (p = 1; Fig. 1C) despite mRNA expression of *nspf-1* during this developmental stage (Harris *et al*., 2019). The non-overlapping mRNA and protein localization patterns are again likely to be a sensitivity constraint of our whole-animal immunostaining procedure to clearly identify subcellular localization patterns. Adult hermaphrodites all showed the same vulva and uterus localization pattern of staining as we observed in mated, control females (Fig. 1C; Fig. S1). On days 1 and 2 of adulthood, 54% (n = 54 of 100) and 41% (n = 61 of 150) of hermaphrodites, respectively, showed staining, which was significantly more than control females (Day 1: χ^2^ = 49.9, df = 1, p < 0.0001; Day 2: χ^2^ = 22.6, df = 1, p < 0.0001). On day 8 of adulthood, 67% (n = 90 of 134) of hermaphrodites showed the same significant staining pattern (χ^2^ = 147.2, df = 6, p < 0.0001; Fig. 1C). Because self-sperm are depleted by day 8 late in adulthood (Hirsh et al. 1976), these data further support the male transfer of NSPF proteins to the uterus. Across all developmental time points, we noted the unexpected absence of signal in the spermathecae, likely due to sensitivity of the whole-worm staining procedure.

To distinguish whether the early adulthood signal of NSPF staining was caused by self-fertilization or mating, we examined a male-less population of His-tagged hermaphrodites on day 1 of adulthood prior to the start of egg laying. We found that 59% (n = 59 of 100) of hermaphrodites showed the same vulva and uterus pattern of staining (Fig. 3D) despite no opportunity for male transfer of NSPF proteins. This frequency was not significantly different from the number of day 1 hermaphrodites raised in the presence of males (p = 0.57). These results suggest that the uterine localization of the NSPF proteins – whether secreted by self-sperm or male sperm – likely reflects their site of activity and not simply a byproduct of the point of transfer during mating.

### NSPF genes provide a weak fitness benefit

Having demonstrated that the NSPF proteins are part of seminal fluid, we sought to discover their relationship with reproductive fitness (Fig. S4). Under a model in which expression of *nspf* genes exert a net deleterious effect on females and males, we would expect the evolutionary loss of *nspf* genes from the genomes of species that evolved primarily selfing hermaphroditism (Yin *et al*., 2018; Cutter *et al*., 2019), with dynamic patterns of gain and loss for the gene family across species given the high divergence among members of the *Caenorhabditis* genus (Konrad *et al*., 2018; Stevens *et al*., 2020). Alternatively, under a model in which NSPF activity is selectively-neutral, we would expect to see a lack of constraint in the sequence evolution and copy number dynamics of this gene family (Kimura, 1968). However, the signatures of conservation of amino acid sequence and synteny indicating purifying selection (Kasimatis *et al*., 2018) are consistent with neither a deleterious nor a neutral model. Instead, they suggest a net fitness benefit mediated by one or both sexes. If the NSPF proteins were neutral or beneficial to male reproductive function but costly to female reproductive function, then we would expect: 1) self-sperm derived NSPF expression to be lost or significantly reduced (Yin *et al*., 2018; Cutter *et al*., 2019), and/or 2) dynamic molecular evolution of the gene family across the phylogeny due to heterogeneity in the net balance of male benefit to female cost. Given the NSPF phylogenetic conservation (Kasimatis *et al*., 2018) as well as conservation of self-sperm derived NSPF at the transcriptional level (Ebbing *et al*., 2018) and protein level (Fig. 3D), we hypothesize that the NSPF proteins benefit reproductive fitness and that this benefit is directed toward both female and male reproduction (Fig. S4).

Previous experiments on male fertility as young adults showed no significant difference in total fecundity between wildtype males and males with all three copies of the NSPF genes deleted (*i.e.,* deletion allele) (Fig. S3A; Kasimatis *et al*., 2018). Using old males (day 5 of adulthood), we repeated these non-competitive, total fecundity assays to determine whether NSPF proteins improved male sperm fertilization ability in a late phase of the life cycle when male fertility is declining overall. Although the mean fecundity of old males with the deletion allele trended lower than wildtype males (*x_del_* = 459; *x_wt_* = 481.9), the effect was not significant (p = 0.85; Fig. S3A). This nominal decrease in fecundity of males with the deletion allele was similar to what was previously seen with young adult males under a competitive male setting, suggesting that NSPF function is potentially both age-independent and sperm competition-independent (Kasimatis et al. 2018; Fig. S3B). Given the experiment size and high reproductive variance, however, a difference between genotypes of 71% would be required to achieve 80% power to detect a significant difference. Nonetheless, the within-generation trends suggest that these proteins are either beneficial or neutral with regards to both male and female fitness, despite the high abundance of the proteins. The null results of these late-life assays coupled with the previous early-life assays (Kasimatis *et al*., 2018) provide no evidence that these proteins are costly to females as the fecundity of females mated with deletion males did not increase in three independent experiments.

We turned to experimental evolution to overcome the power limitations of within-generation experimental design and test whether the NSPF genes are male-beneficial or male-neutral. Specifically, we competed the wildtype allele against the deletion allele in an obligate outcrossing *C. elegans* population by evolving ten replicate populations of 1,000 individuals from a base population that was heterozygous at the NSPF locus in an otherwise homogeneous genetic background. Between generations 11 and 20, we documented a significant overall decrease in the frequency of the deletion allele across replicates (GLM Model 1: z-value = −2.9, p = 0.003) with the deletion allele clearly decreasing in frequency by at least 4% in four of the ten replicates and remaining approximately the same in two replicates (≤2% change). Between replicate effects accounted for 27% of the variance observed. When including the sequencing control samples as a technical estimate of the allele frequencies at generation 1, no significant allele frequency change is observed over time, suggesting a more neutral fitness effect of the NSPF protein (Fig. 4; Table S3). These results support the within-generation fecundity trends to indicate that any effect of males transferring NSPF protein to females does not impose a large fitness cost to females. Instead, under these conditions, we estimate the wildtype allele to confer approximately a 0.1% fitness advantage over the deletion allele (Table S3). Together, these proteins appear to provide an overall weak net fitness effect that is consistent with being beneficial or neutral to females and males (Fig. S4).

**Figure 4.**
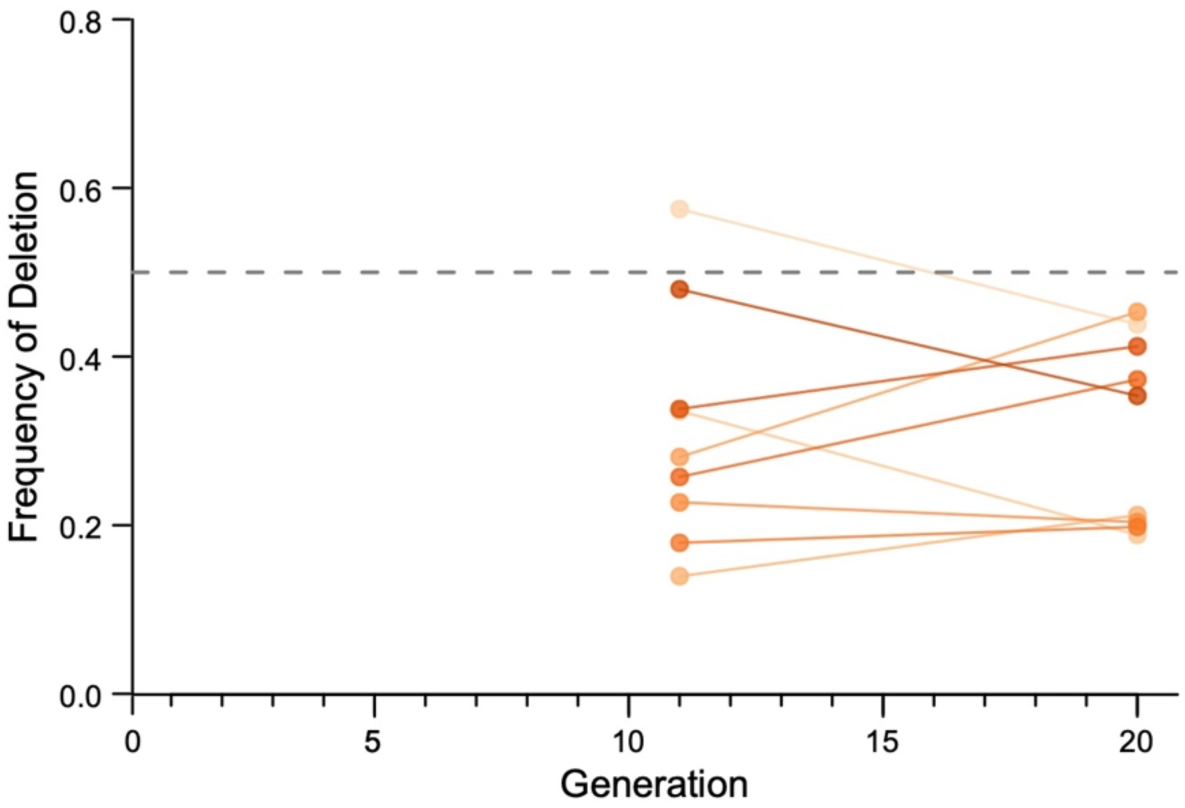
Changes in the frequency of the deletion allele over time estimated using amplicon sequencing data. All ten replicates started with 50% deletion allele (dashed line). The orange points represent the frequency of the deletion allele in each replicate population empirically estimated from amplicon sequence data collected at generations 11 and 20. Statistical testing of the empirical observations used a general linear model (Table S3). There was a significant decrease in the frequency of the deletion allele between generations 11 and 20 (GLM Model 1: s = 0.001, z-value = −2.98, p < 0.01).

## DISCUSSION

Seminal fluid is a complex bioactive mixture whose components are implicated in a variety of reproductive, behavioral and physiological functions and thus likely evolves in response to multiple selective pressures. We took an integrative approach to investigate the highly abundant MO-associated NSPF proteins. We confirmed that the NSPF genes are indeed a component of seminal fluid that is transferred to females, which demonstrates that MO fusion provides a novel source of seminal fluid proteins. Additionally, the localization of NSPF proteins (*i.e.,* the functional output) differs from that of mRNA transcripts, which is consistent with ejaculate proteins and not sperm-specific proteins. Finally, using within- and across-generation experiments, we quantified the small, but significant contribution made to reproductive fitness by the presence of NSPF proteins, most consistent with a positive fitness effect in males and either a positive or neutral fitness effect in females (Fig. S4). Together, this work demonstrates the power of *Caenorhabditis* to expand our perspective of seminal fluid protein function and to disentangle beneficial versus antagonistic effects of seminal fluid proteins.

The localization of NSPF protein to the spicule (Fig. 2) and uterus (Fig. 3) was unexpected given that the NSPFs were previously characterized as being associated with sperm MOs (Kasimatis *et al*., 2018) and that mRNA expression data indicated that all three *nspf* genes are expressed in the gonad only (Ebbing *et al*., 2018). Staining of dissected testes provided the necessary sensitivity to confirm that NSPF proteins localize to spermatid MOs. Our whole-worm staining approach did, however, identify additional regions of localization that may result from the anatomical difference in the location of inactivated versus activated sperm. The activation process occurs as sperm pass from a male to a female/hermaphrodite while the spicules are everted in the vulva, thus creating a direct link between MO fusion and the spicules. This overlap in timing, however, does not explain the spicule signal seen in L4 larval males, which warrants further work. In hermaphrodites, the parsimonious hypothesis is that NSPF proteins passively diffuse from the spermathecae, where activated sperm reside, to the uterus. The location of NSPF proteins within the uterus is similar to other proteins that males transfer, which originate in newly identified extracellular vesicles (Nikonorova *et al*., 2022). While the NSPFs do not appear in these vesicles, it is possible that this alternative vesicle type also contributes to seminal fluid proteins, supporting our finding that nematodes employ a distinct mechanism for secreting seminal fluid proteins than either mammals or *Drosophila* (McGraw *et al*., 2015).

We capitalized on the ability to manipulate mating system to show that NSPF production is not only retained during the lineage transition to self-fertilization, but that the proteins localize to the same uterus region in unmated hermaphrodites as in mated females (Fig. 3). This pattern reinforces the pattern of conserved molecular evolution of the *nspf* gene sequences (Kasimatis *et al*., 2018) and implies that the reproductive function of the sperm-dependent NSPF proteins within the uterus is mating-independent. Importantly, the conserved expression during spermatogenesis in L4 hermaphrodites relative to males of the ancestral gonochoristic species supports the notion that these proteins are not involved in ongoing sexual conflict, as there would be strong selection favoring their loss as seen in many other male reproductive traits (Thomas *et al*., 2012; Fierst *et al*., 2015; Yin *et al*., 2018). Instead, expression of *nspf* genes appear beneficial to reproductive fitness, if weakly, as supported by fitness assays. If their net advantage resulted from a detriment to females that was offset by a male fitness benefit, then we would expect females mated with deletion males to trend toward higher reproductive success than females mated with wildtype males. However, our experiments support the opposite of such a scenario, suggesting NSPF promotes sexually-concordant reproductive fitness.

A common challenge in evolutionary biology is to detect the fitness effects of variants at a single gene (Barrett & Hoekstra, 2011). Our experimental evolution design to compete the wildtype and deletion alleles in a highly-replicated fashion allowed for the potential to compound small effect sizes over tens of generations. The nematode model system allows for this approach to be done at a larger scale and over a longer time periods than similar studies (Wong & Rundle, 2013), which, when coupled with genomics, provides greater power to detect small or fluctuating selection coefficients that are most likely to underlie allelic effects in standing genetic variation. The overall trend across replicates was a slight decrease in the frequency of the deletion allele (Fig. 4). The estimated selection coefficient, despite being small, would be powerful enough to prevent *nspf* gene loss over time in nature. Such a small fitness effect is surprising given the molecular evolutionary history of these genes, but supports the “mild phenotype” observed in many classic genetics knockout/knockdown studies (Sugimoto, 2004; Bell, 2010). The frequency of the deletion allele did not decrease consistently in every replicate at each sampling time point, however, which likely reflected a combination of variance and stochasticity during mating along with sampling error and the relatively small selection coefficient. Additionally, the ability of females to mate with multiple males created the opportunity for wildtype males to compensate for any benefits a female lost by mating with a deletion male. Such compensation would dampen the effect of selection against the deletion allele and potentially contribute to the fluctuating frequency of the deletion allele. Overall, this approach of allelic competition in experimental evolution (Gray & Cutter, 2014; Teotonio *et al*., 2017) may represent a promising avenue for future seminal fluid protein studies, and for genetic variants of other phenotypes, where small fitness effect sizes have proved challenging to characterize.

Our data motivate multiple hypotheses for future testing to narrow down the specific beneficial function of the NSPF proteins. First, the NSPF proteins could interact with a male-female signaling network that promotes egg laying. While the interacting partner for NSPF proteins remains unknown, it is possible that NSPFs bind to muscle proteins that form the uterus walls and are involved in egg laying behavior. Under this hypothesis, localization to the spicules is puzzling. However, previous work has shown the females pressurize the gonad to push some of the ejaculate mass and spicules out of the vulva (Barker, 1994; Kleemann & Basolo, 2007). This process could create a residue on the spicule cuticle as a byproduct, though we cannot rule out a functional relationship between the NSPF proteins and spicule cuticle.

Alternatively, the NSPF proteins could impact the uterine environment. Specifically, seminal fluid of some species contains anti-microbial factors (Samakovlis *et al*., 1991; Gao & Zhu, 2016). Given the density of protein right above the vulva, one hypothesis is that the NSPF proteins play an anti-microbial role to protect the uterus from bacterial infection. Previous work has shown that the uterus can be exposed to pathogenic infection from the external environment during mating and egg laying (Muir & Tan, 2008). The male spicules are presumably also exposed during eversion, and thus any anti-microbial activity of the NSPF residue on the spicules might serve to prevent bacterial infection in males when the spicules retract. Biochemically, the alpha-helix structure of NSPF proteins lends plausibility to this hypothesis, as such secondary structures can confer generic bacterial lysing properties (Huan *et al*., 2020). Future work testing the fitness effects of the NSPF deletion allele in worms raised in environments with more pathogenic bacteria is warranted.

Our work demonstrates the complex but subtle dynamics of seminal fluid proteins uncovered when using integrative approaches. We also showed how *Caenorhabditis* nematodes are an excellent system for expanding our knowledge of post-insemination interactions not only for their technical benefits, but also for their novel sperm components and potential for distinguishing sexual conflict from sexual cooperation by making mating system contrasts between male-derived sperm and self-sperm interactions. Connecting the layers of the genotype-phenotype-fitness map for seminal fluid proteins will help us uncover the genetics of fertility and the evolution of reproductive success.

## Acknowledgements

We thank the Phillips lab for sharing plasmids pMS4 and pZCS16 and strain PX814. We thank Zachary Stevenson for transgenic advice, Jack Hu and Mathias Renaud for immunostaining advice, and Stephen Banse, Eric Haag and the Cutter and Claycomb labs for their helpful discussion.

## Author Contributions

KRK, ADC, and LR devised the project. KRK created strains VX194 and VX300. KRK performed the fecundity assay and experimental evolution. CR performed the immunostaining assays with assistance from KRK. KRK analyzed the data. KRK wrote the manuscript with the support of the other authors.

## Data accessibility

The oligonucleotides used in this study are available as Table S1. The immunostaining data are available in File S2 and the complete protocol is available in File S1. The fertility data are available in File S3. The raw amplicon data are available at NCBI SRA under accession number PRJNA1009866. The processed amplicon reads are available in File S4. All scripts are available via the Cutter Lab GitHub repository NSPF (https://github.com/Cutterlab/NSPF). Worm strains JK574 and VX300 are available from the *Caenorhabditis* Genetics Center. Strains PX814 and VX194 are available from the Cutter lab upon request.

The authors declare no conflicts of interest.

## Supplemental Files

**Fig S1. Immunostaining quantification development. A)** Count data showing the percent of control males (JK574, triangle) and His-tagged males (VX300, circle) with fluorescence in each anatomical region during larval stage 4 (L4, orange), day 1 of adulthood (D1, green), day 2 of adulthood (D2, blue), and day 8 of adulthood (D8, pink). **C)** Count data showing the percent of control females (JK574, triangle) and His-tagged hermaphrodites (VX300, circle) with fluorescence in each anatomical region during larval stage 4 (L4, orange), day 1 of adulthood (D1, green), day 2 of adulthood (D2, blue), and day 8 of adulthood (D8, pink).

**Fig S2. NSPF-1 localizes the MOs of spermatids in adult males. A)** Protein (green) localized in a punctate pattern around the nucleus (blue) of spermatids in the dissected testes of adult His-tagged males. **B)** Protein localization within a single spermatid (outlined) of an adult His-tagged male. **C)** Protein localization within multiple spermatids (arrows) of an adult His-tagged male.

**Fig S3. Single male fecundity assays. A)** The total fecundity of wildtype males with the NSPF genes (gray) and deletion males with the NSPF genes removed (orange) at day 1 of adulthood and day 5 of adulthood. There was no significant difference in reproductive success between male types. Day 1 data were replotted from Kasimatis et al. (2018). **B)** The proportion of progeny coming from wildtype males (gray) or deletion males (orange) versus a wildtype competitor male. There was no significant difference in sperm competitive ability between male types (Kasimatis et al. 2018). Competitive fecundity success data were replotted from Kasimatis et al. (2018).

**Fig S4. Summary of evidence for the fitness effect of NSPF in each sex.** We present the expected pattern of sequence evolution, gene expression, immunostaining, sperm biology, and allele frequency for each combination of NSPF effects on males and females. The lines of evidence supporting or contrary to that combination of effects is then given. The evidence most strongly supports a fitness model of male beneficial and female beneficial or neutral.

**Table S1. Primers used in this study.**

**Table S2. Strains used in this study.**

**Table S3. Model comparison for amplicon sequencing data.**

**File S1. Complete immunostaining protocols and buffer recipes.**

**File S2. Immunostaining data.**

**File S3. Late life fecundity data.**

**File S4. Processed amplicon reads.**

## References

Albritton, S.E., Kranz, A.-L., Rao, P., Kramer, M., Dieterich, C. & Ercan, S. 2014. Sex-biased gene expression and evolution of the x chromosome in nematodes. Genetics 197: 865–883.

Andersen, E.C., Gerke, J.P., Shapiro, J.A., Crissman, J.R., Ghosh, R., Bloom, J.S., et al. 2012. Chromosome-scale selective sweeps shape *Caenorhabditis elegans* genomic diversity. Nat Rev Genet 44: 285–290.

Baldi, C., Cho, S. & Ellis, R.E. 2009. Mutations in two independent pathways are sufficient to create hermaphroditic nematodes. Science 326: 1002–1005.

Barker, D.M. 1994. Copulatory plugs and paternity assurance in the nematode *Caenorhabditis elegans*. Animal Behaviour 48: 147–156.

Barrett, R.D.H. & Hoekstra, H.E. 2011. Molecular spandrels: tests of adaptation at the genetic level. Nat Rev Genet 12: 767–780.

Begun, D.J., Whitley, P., Todd, B.L., Waldrip-Dail, H.M. & Clark, A.G. 2000. Molecular population genetics of male accessory gland proteins in *Drosophila*. Genetics 156: 1879– 1888.

Bell, G. 2010. Experimental genomics of fitness in yeast. Proc. R. Soc. B: Biol. Sci. 277: 1459– 1467.

Beydoun, S., Choi, H.S., Dela-Cruz, G., Kruempel, J., Huang, S., Bazopoulou, D., et al. 2021. An alternative food source for metabolism and longevity studies in *Caenorhabditis elegans*. Commun Biol 4: 258–9.

Brenner, S. 1974. The genetics of *Caenorhabditis elegans*. Genetics 77: 71–94.

Chapman, T. 2011. Seminal fluid-mediated fitness traits in *Drosophila*. Heredity 87: 511–521.

Clark, N.L., Aagaard, J.E. & Swanson, W.J. 2006. Evolution of reproductive proteins from animals and plants. Reproduction 131: 11–22.

Cutter, A.D., Morran, L.T. & Phillips, P.C. 2019. Males, Outcrossing, and Sexual Selection in Caenorhabditis Nematodes. Genetics 213: 27–57.

Dapper, A.L. & Wade, M.J. 2016. The evolution of sperm competition genes: The effect of mating system on levels of genetic variation within and between species. Evolution 70: 502– 511.

Dickinson, D.J., Pani, A.M., Heppert, J.K., Higgins, C.D. & 2015. 2015. Streamlined genome engineering with a self-excising drug selection cassette. Genetics 200: 1035–1049.

Ebbing, A., Vértesy, Á., Betist, M.C., Spanjaard, B., Junker, J.P., Berezikov, E., et al. 2018. Spatial transcriptomics of *C. elegans* males and hermaphrodites identifies sex-specific differences in gene expression patterns. Developmental Cell 47: 801–813.

Ellis, R.E. & Stanfield, G.M. 2014. The regulation of spermatogenesis and sperm function in nematodes. Sem Cell Dev Biol 29: 17–30.

Emmons, S.W. 2005. Male development. WormBook 1–22.

Félix, M.-A. & Duveau, F. 2012. Population dynamics and habitat sharing of natural populations of *Caenorhabditis elegans* and *C. briggsae*. BMC Biol. 10: 59.

Fierst, J.L., Willis, J.H., Thomas, C.G., Wang, W., Reynolds, R.M., Ahearne, T.E., et al. 2015. Reproductive mode and the evolution of genome size and structure in *Caenorhabditis* nematodes. PLoS Genetics 11: e1005323.

Gao, B. & Zhu, S. 2016. The drosomycin multigene family: three-disulfide variants from Drosophila takahashii possess antibacterial activity. Nat Rev Genet 6: 32175.

Gray, J.C. & Cutter, A.D. 2014. Mainstreaming *Caenorhabditis elegans* in experimental evolution. Proc Royal Soc B Biological Sci 281: 20133055.

Hansen, J.M., Chavez, D.R. & Stanfield, G.M. 2015. COMP-1 promotes competitive advantage of nematode sperm. eLife 1–26.

Harris, T.W., Arnaboldi, V., Cain, S., Chan, J., Chen, W.J., Cho, J., et al. 2019. WormBase: a modern model organism information resource. Nucleic Acids Research 29: 82.

Hopkins, B.R. & Perry, J.C. 2022. The evolution of sex peptide: sexual conflict, cooperation, and coevolution. Biological Reviews 97: 1426–1448.

Huan, Y., Kong, Q., Mou, H. & Yi, H. 2020. Antimicrobial peptides: classification, design, application and research progress in multiple fields. Front Microbiol 11: 582779.

Kasimatis, K.R., Moerdyk-Schauwecker, M.J., Timmermeyer, N. & Phillips, P.C. 2018. Proteomic and evolutionary analyses of sperm activation identify uncharacterized genes in *Caenorhabditis* nematodes. BMC Genomics 19: 593.

Kenyon, C. 1988. The Nematode *Caenorhabditis elegans*. Science 240: 1448–1453.

Kimura, M. 1968. Evolutionary rate at the molecular level. Nature 217: 624–626.

Kleemann, G.A. & Basolo, A.L. 2007. Facultative decrease in mating resistance in hermaphroditic *Caenorhabditis elegans* with self-sperm depletion. Anim Behav 74: 1339– 1347.

Konrad, A., Flibotte, S., Taylor, J., Waterston, R.H., Moerman, D.G., Bergthorsson, U., et al. 2018. Mutational and transcriptional landscape of spontaneous gene duplications and deletions in *Caenorhabditis elegans*. Proc. Natl. Acad. Sci. 115: 7386–7391.

LaMunyon, C.W. & Ward, S. 1998. Larger sperm outcompete smaller sperm in the nematode *Caenorhabditis elegans*. Proc R Soc Lond B 265: 1997–2002.

McDonough, C.E., Whittington, E., Pitnick, S. & Dorus, S. 2016. Proteomics of reproductive systems: Towards a molecular understanding of postmating, prezygotic reproductive barriers. J. Proteomics 135: 26–37.

McGraw, L.A., Suarez, S.S. & Wolfner, M.F. 2015. On a matter of seminal importance. Bioessays 37: 142–147.

Muir, R.E. & Tan, M.-W. 2008. Virulence of *Leucobacter chromiireducens* subsp. *solipictus* to *Caenorhabditis elegans*: characterization of a novel host-pathogen interaction. Appl Environ Microbiol 74: 4185–4198.

Nikonorova, I.A., Wang, J., Cope, A.L., Tilton, P.E., Power, K.M., Walsh, J.D., et al. 2022. Isolation, profiling, and tracking of extracellular vesicle cargo in *Caenorhabditis elegans*. Curr. Biol. 32: 1–13.

Perry, J.C., Sirot, L. & Wigby, S. 2013. The seminal symphony: how to compose an ejaculate. Trends Ecol Evol 28: 414–422.

Poiani, A. 2006. Complexity of seminal fluid: a review. Behav Ecol Sociobiol 60: 289–310.

Richaud, A., Zhang, G., Lee, D., Lee, J. & Félix, M.-A. 2018. The local coexistence pattern of selfing genotypes in *Caenorhabditis elegans* natural metapopulations. Genetics 208: 807– 821.

Rowe, L., Chenoweth, S.F. & Agrawal, A.F. 2018. The genomics of sexual conflict. Am Nat 192: 274–286.

Samakovlis, C., Kylsten, P., Kimbrell, D.A., Engström, A. & Hultmark, D. 1991. The andropin gene and its product, a male-specific antibacterial peptide in *Drosophila melanogaster*. The EMBO Journal 10: 163–169.

Shakes, D.C., Wu, J., Sadler, P.L., LaPrade, K., Moore, L.L., Noritake, A., et al. 2009. Spermatogenesis-specific features of the meiotic program in *Caenorhabditis elegans*. Plos Genet 5: e1000611.

Smith, J.R. & Stanfield, G.M. 2011. TRY-5 Is a sperm-activating protease in *Caenorhabditis elegans* seminal fluid. PLoS Genetics 7: e1002375.

Stevens, L., Rooke, S., Falzon, L.C., Machuka, E.M., Momanyi, K., Murungi, M.K., et al. 2020. The genome of *Caenorhabditis bovis*. Curr. Biol. 30: 1023–1031.e4.

Sugimoto, A. 2004. High-throughput RNAi in *Caenorhabditis elegans*: genome-wide screens and functional genomics. Differentiation 72: 81–91.

Swanson, W.J. & Vacquier, V.D. 2002. Reproductive protein evolution. Annu. Rev. Ecol. Syst. 33: 161–179.

Team, R.C. 2020. R: A language and environment for statistical computing. Foundation for Statistical Computing.

Teotonio, H., Estes, S., Phillips, P.C. & Baer, C.F. 2017. Experimental evolution with *Caenorhabditis* nematodes. Genetics 206: 691–716.

Thomas, C.G., Li, R., Smith, H.E., Woodruff, G.C., Oliver, B. & Haag, E.S. 2012. Simplification and desexualization of gene expression in self-fertile nematodes. Current Biology 22: 2167– 2172.

Ting, J.J., Woodruff, G.C., Leung, G., Shin, N.-R., Cutter, A.D. & Haag, E.S. 2014. Intense sperm-mediated sexual conflict promotes reproductive isolation in *Caenorhabditis* nematodes. PLoS Biol 12: e1001915–14.

Wilburn, D.B. & Swanson, W.J. 2016. From molecules to mating: Rapid evolution and biochemical studies of reproductive proteins. J. Proteomics 135: 12–25.

Wong, A. & Rundle, H. 2013. Selection on the *Drosophila* seminal fluid protein Acp62F. Ecol Evol 3: 1942–1950.

Yin, D., Schwarz, E.M., Thomas, C.G., Felde, R.L., Korf, I.F., Cutter, A.D., et al. 2018. Rapid genome shrinkage in a self-fertile nematode reveals sperm competition proteins. Science 359: 55–61.

