## Supplementary figures and images for "A novel sperm-derived seminal fluid protein in *Caenorhabditis* nematodes"

### Figure S1

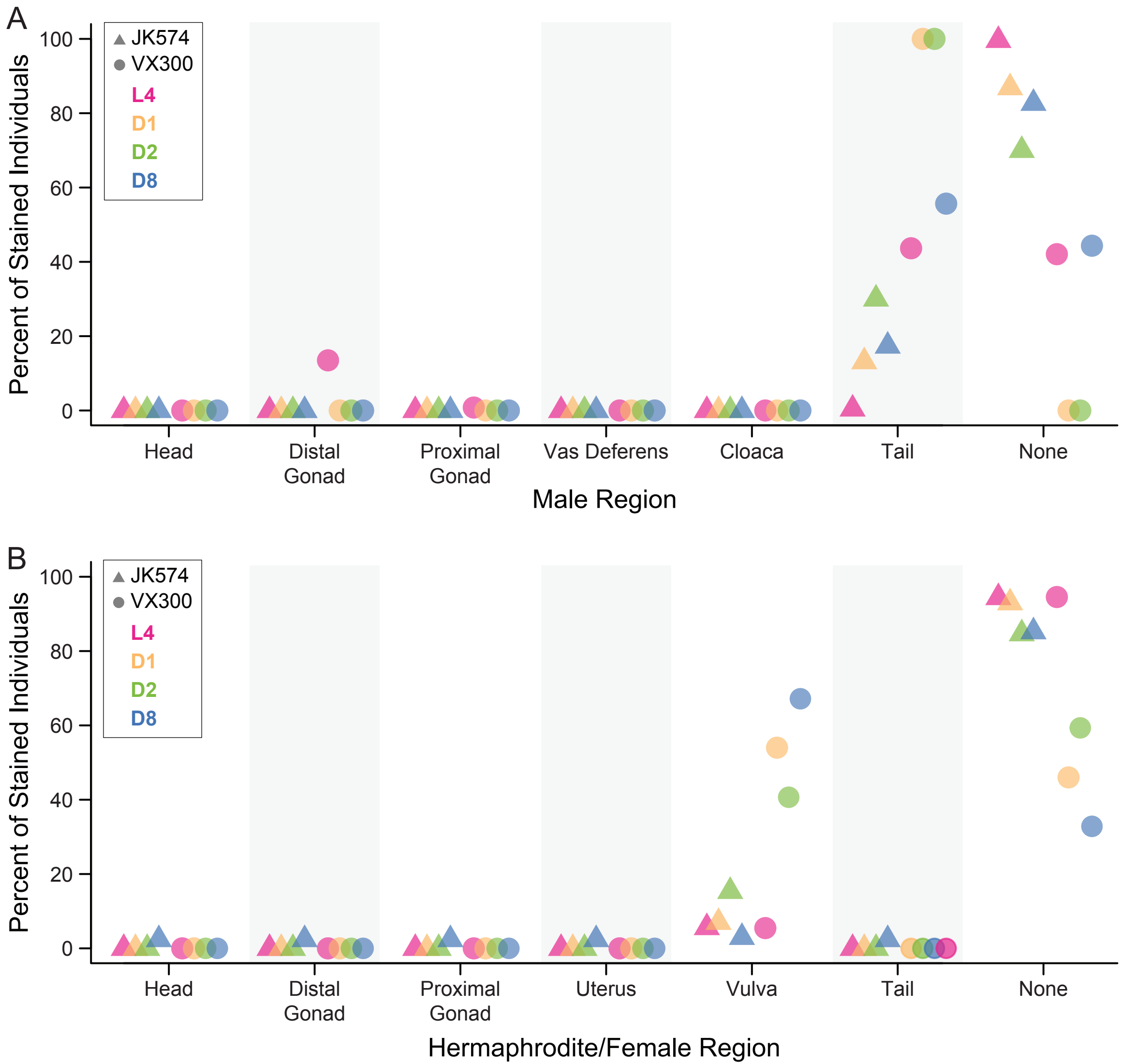

### Figure S2

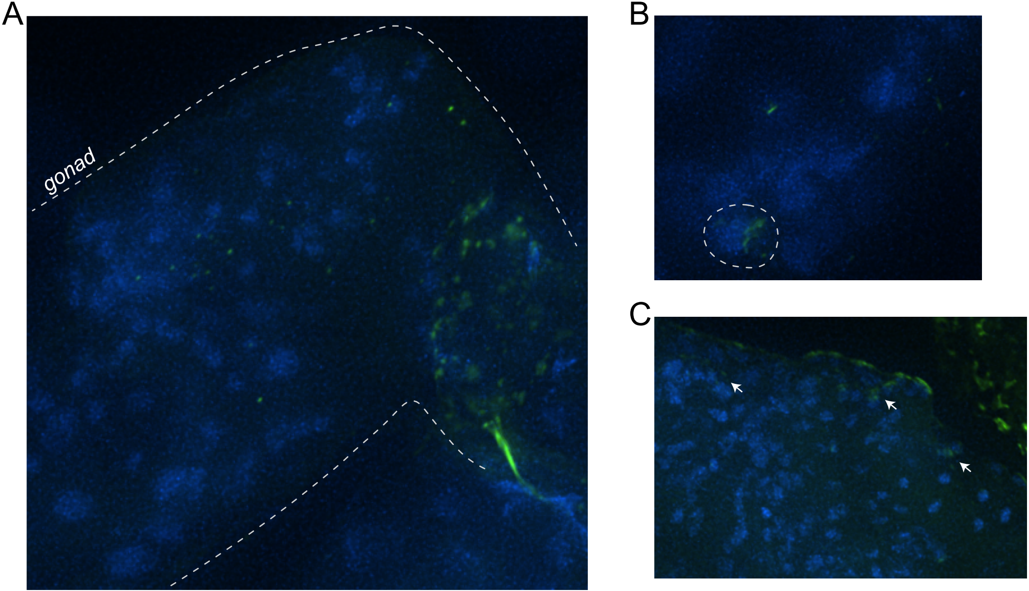

### Figure S3

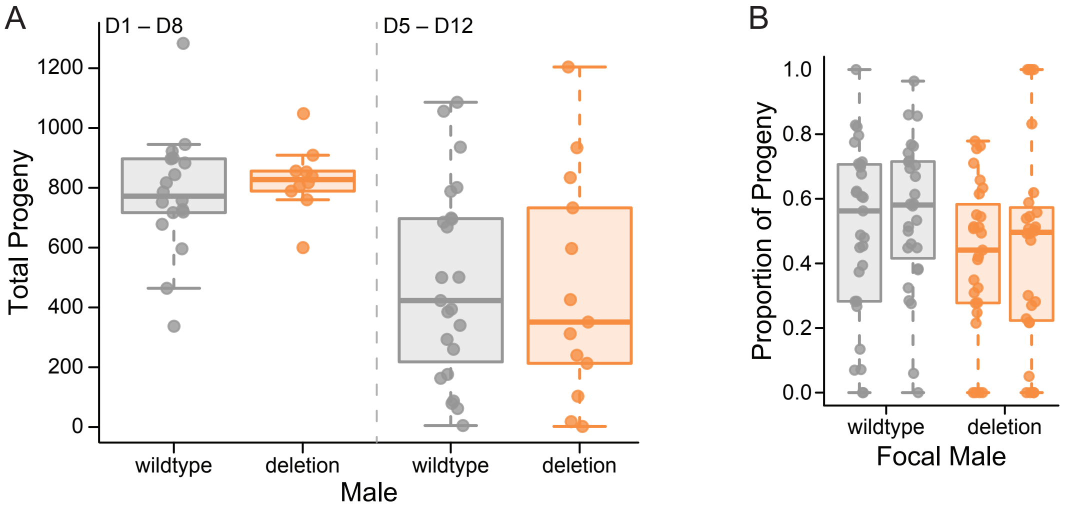
