## Supplementary material for "A novel sperm-derived seminal fluid protein in *Caenorhabditis* nematodes": Figure S4

| | Male Effect × Female Effect[1] | | | | | | | | |
|---|---|---|---|---|---|---|---|---|---|
| | **B × B** | **B × N** | **B × C** | **N × B** | **N × N** | **N × C** | **C × B** | **C × N** | **C × C** |
| **Expectation** Sequence Evolution[2] | Purifying selection | Purifying selection | Positive selection in outcrossers; Net directional selection for gene loss in hermaphrodites | Purifying selection | Molecular evolution and/or gene loss via genetic drift | Directional selection for gene loss | Positive selection | Directional selection for gene loss | Directional selection for gene loss |
| Male Expression[3] | Present | Present | Present | Possible | Possible | Possible | Absent | Absent | Absent |
| Hermaphrodite Expression[3] | Present | Possible | Absent (after long evo. divergence time[7-9]) | Present | Possible | Absent | Present | Possible | Absent |
| Male Immunostaining[4] | Present | Present | Present | Possible | Possible | Possible | Absent | Absent | Absent |
| Hermaphrodite Immunostaining[4] | Present | Possible | Absent (after long evo. divergence time[7-9]) | Present | Possible | Absent | Present | Possible | Absent |
| Packaged into Male Sperm[5] | Present | Present | Present | Possible | Possible | Possible | Absent | Absent | Absent |
| Exp. Evo. Allele Frequency[6] | Directional selection favoring WT allele | Directional selection favoring WT allele | *not addressed by this experimental design* | Directional selection favoring WT allele | Genetic drift; high replicate variance | Directional selection favoring DEL allele | *not addressed by this experimental design* | Directional selection favoring DEL allele | Directional selection favoring DEL allele |
| **Supporting Lines of Evidence** | 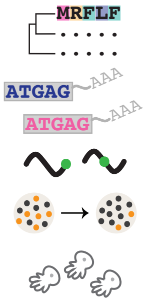 | 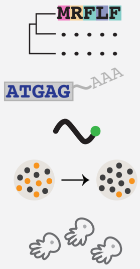 | 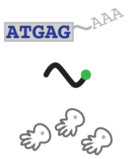 | 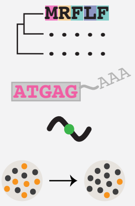 | | | 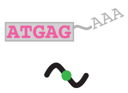 | | |
| **Contrary Lines of Evidence** | | | 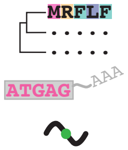 | | 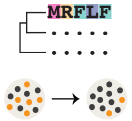 | 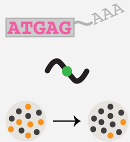 | 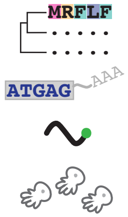 | 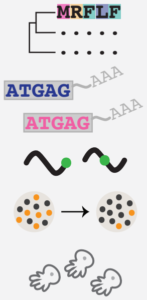 | 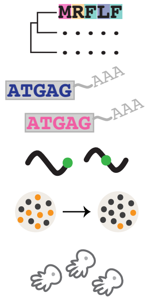 |
| **Conclusion** | **Strong evidence** | **Strong evidence** | Weak evidence | Moderate evidence | No evidence | No evidence | Weak evidence | No evidence | No evidence |

[1]Effect of NSPF genes are beneficial (B), neutral (N), or costly (C) for male function and female function; [2]Molecular evolution signatures from Kasimatis et al. (2018); [3]Expression data from Albritton et al. (2014); [4]Male immunostaining data from this paper; [5]Sperm localization data from Kasimatis et al. (2018) and this paper; [6]Experimental evolution data from this paper; [7]Fierst et al. (2015); [8]Thomas et al. (2012); [9]Yin et al. (2018)
