## Supplementary material for "A novel sperm-derived seminal fluid protein in *Caenorhabditis* nematodes": File S1

Supplementary Methods: Antibody Staining of Formaldehyde-Fixed Worms

modified from Ruvkun and Finney along with and Beydoun et al. (2021)

Note: This protocol works for all stages except dauers and hypochlorite-treated eggs.

**At least 4 days in advance**

*Make plates*

A. Pour NGM-agar plates with carbenicillin (final concentration 100ng/mL).

*Make dead food following Beydoun et al. (2021)*

B. Formaldehyde kill a fresh 50mL culture of OP50 *E. coli* by adding 2.5mL of *10% formaldehyde* (final concentration 0.5%) and shaking (170 rpm) at 37C for 1hr.
C. Divide the culture into two 50mL conical tubes and centrifuge at 3000rcf for 15min to pellet the cells.
D. Remove the supernatant. Add ~15mL of fresh *LB+Triton* (0.5%) and centrifuge for 8min. Repeat this step for a total of five washes.
E. After the final wash, pellet the cells and add 5-6mL of *LB*. Distribute to NGM+carbenicillin plates using a Pasteur pipette. (In our hands, worms grew more effectively if the dead OP50 lawn was not spread into a thin layer, but rather left as a large “spot” in the center of the plate.)

*Stage worms*

F. Hatch-off worms (or do an egg lay) and stage appropriately on NGM+carbenicillin plates.

**Day 1: Fixation**

*Prepare worms*

1. Wash worms into a falcon tube with *M9*. Spin at 1400 rcf for 2min and remove supernatant. Add 5mL *M9* + *Triton* (1%) and repeat the spin.
2. Add 5mL *M9* + *Triton* (1%) and gently rock worms (e.g., on a nutator) at room temperature* for 20min. Spin down and remove the supernatant.
3. Repeat step 2 two more times for a total of three soaks.
4. Spin down and remove the supernatant. Add 5mL of *autoclaved MilliQ water*. Spin down and remove the supernatant. Repeat the spin with water two more times.
5. After the final spin, transfer the pelleted worms to an Eppendorf tube using a glass Pasteur pipette. Spin worms at 400 rcf for 2min and remove as much supernatant as possible. Worms are now ready for fixation.

*Prepare Fixative and Blocking Buffer*

6. While the worms are soaking, dissolve 0.025g *skim milk* in 5mL *Blocking Buffer*.
7. While the worms are in the final soak, thaw a 10% formaldehyde aliquot. (Also remove an aliquot of DTT and keep on ice.)
8. Vortex the formaldehyde solution until it is mostly clear. Filter into a fresh tube using a 0.22um pore syringe filter and keep on ice.
9. When the worms are ready, prepare *fresh fixative* (see “Recipes”).

*Formaldehyde-fix worms*

10. Fixation: Add 1,250uL of *fresh fixative* per tube. Mix well by gently inverting.
11. Immerse the tube in dry ice + ethanol (or liquid nitrogen) to freeze. Thaw the tube contents on ice and incubate on ice with occasional agitation for 30min.
12. Spin worms 400 rcf for 2min and remove the supernatant.
13. Reduction: Wash worms twice with 2mL *1x Tris-Triton Buffer*. Resuspend worms in 1mL *1x Tris-Triton Buffer* + 10uL *β-mercaptoethanol* (1%). Incubate at 37C for 1.5hr with gentle shaking (140 rpm). After this point, the worms are fragile and should not be spun hard.
14. Spin worms at 200 rcf for 2min and remove the supernatant.
15. Resuspend in 1mL *1x Borate Buffer*. Spin worms down and remove the supernatant.
16. Resuspend in 500uL *1x Borate Buffer* + 500uL *20mM DTT* (final concentration 10mM DTT). Rock gently at room temperature for 15min.
17. Spin worms down and remove the supernatant.
18. Oxidation: Resuspend in 1mL *1x Borate Buffer* + 3uL *H_2_O_2_* (0.3%). Wrap the lids with parafilm and rock gently at room temperature for 15min.
19. Spin worms down and remove the supernatant. Resuspend in 1mL *1x Borate Buffer* and repeat spin.
20. Blocking: Resuspend worms in 1mL *Blocking Buffer* + *Skim Milk* (0.5%). Rock gently at room temperature for 30min.
21. Spin worms down and remove the supernatant.
22. Resuspend worms in 1mL *Antibody-B Buffer* and rock gently at room temperature for 15min. Spin down at 400 rcf for 2 min and remove the supernatant. Resuspend in 1mL *Antibody-A Buffer*. At this point worms can be stored at 4C.

**Day 2: Staining**

*Staining*

23. Transfer a 100uL aliquot of fixed worms to an Eppendorf tube for staining. Add an appropriate dilution of antibody and *Antibody-B Buffer* for a total volume of 500uL. Incubate at room temperature for 1hr and rock gently (covered in foil).
24. Spin worms down at 400 rcf for 2min and remove the supernatant.
25. Resuspend in 1mL *Antibody-B Buffer* and repeat spin.
26. Resuspend in 1mL *Antibody-B Buffer* and rock gently at room temperature for 30min (in foil). Spin worms down at 400 rcf for 2min and remove the supernatant. Repeat 3 times for a total of four 30min soaks.

*Co-Staining*

27. Resuspend in 990uL *Antibody-A Buffer* + 10uL DAPI. Incubate at 30C for 30min (in foil).
28. Spin worms and remove the supernatant. Resuspend in 1mL *Antibody-B Buffer* and repeat the spin. Remove the supernatant and resuspend in 1mL *Antibody-A Buffer*. Spin down worms and remove most of the supernatant.
29. Worms are now ready for imaging.

*Note: The room temperature incubation times given throughout “Day 1: Fixation” and “Day 2: Staining” are appropriate for workspaces ranging between 20-22C. If the workspace temperature is outside this range, then we suggest moving all room temperature incubations into a 20C incubator.

**References**

Beydoun S, *et. al.* 2021. An alternative food source for metabolism and longevity studies in *Caenorhabditis elegans*. Commun Biol 4:258.

Ruvkun G and Finney M. “Antibody Staining of Formaldehyde-fixed Worms” https://www.wormatlas.org/antibodystaining.htm.

Supplementary Methods: Antibody Staining of Dissected Gonads

modified from Shakes et al. (2009)

**Day 1** (time required ~5hr)

*Prepare worms*

1. Pick 120 hermaphrodites and 120 males into *separate* low bind 1.5mL Eppendorf tube with 500uL *M9+Triton* (1%). Spin at 400 rcf for 2min and remove supernatant (~450uL).
2. Add 450uL *M9+Triton* (1%) and repeat the spin.
3. Add 450uL *M9+Triton* (1%) and gently shake worms for 30min.
4. Spin worms down and remove the supernatant. Add 147.5uL *M9+Triton* (1%) and 1.5uL NaN_3_.

*Dissect Gonads*

5. Label two ColorFrost Plus slide as “males” and another two as “hermaphrodites”.
6. Pipette 50uL of worms on each of two depression slides. Decapitate worms just below the pharynx and above the gonad using two 30 gauge needles in a scissor motion.
7. When all worms have been decapitated, gently pipette worms onto each of the appropriate ColorFrost Plus sides. Divide all worms between the 2 slides (per sex).

*Fixation and blocking*

8. Freeze-crack: Place the slides on dry ice for 30min. While the slides are on dry ice, prepare two small, labeled containers^#^ of methanol and acetone and place at 4C.
9. Fixation: Immerse the slides in 30mL *95% methanol* for 10min. Remove the slides and place them in 30mL *100% acetone* for 5min. Remove the slides and briefly let them air dry.
10. Place a Petri dish under the microscope to catch buffer. Hold each slide on a slight angle and using Pasteur pipette rinse the slide with *1xPBS + 0.25% Triton*. Distribute the liquid above where the worms are (not directly on top of them) so it flows over the worms. While rinsing look through the scope and make sure that the worms are not washing off.
11. Blocking: Rinse the slides with *Blocking Buffer*.

*Staining*

12. Make a 30mL solution of an appropriate dilution of antibody + *Antibody-A Buffer*. Gently mix and pour into the labeled staining container^#^. Immerse the slides and soak for 1hr at 20C covered in foil.
13. Rinse the slides with *Antibody-A Buffer*. Repeat the rinse two more times.

*Co-staining*

14. Make a 30mL solution of *Antibody-A Buffer* + DAPI. Gently mix and pour into the labeled staining container^#^. Immerse the slides and incubate at 30C for 30min covered in foil.
15. Remove the slides and rinse with *Antibody-A Buffer*. Repeat the rinse.
16. Gonads are now ready for imaging.

^#^Note: Large (150mm diameter) glass petri dishes work well as containers for the various soaking steps. In our hands, 30mL of buffer was the appropriate volume to cover four slides.

**References**

Shakes, D. C. *et al.* 2009. Spermatogenesis-Specific Features of the Meiotic Program in *Caenorhabditis elegans*. Plos Genet 5:e100061.

Recipes

Fixation Solution

1.0mL 2x MRWB

200uL 10% formaldehyde (1% final)

800uL MilliQ H_2_O

2x Modified Ruvkun’s Witches Brew (MRWB)

800uL 2M KCl (160 mM final)

80uL 5M NaCl (40mM final)

2.0mL 0.1M EGTA (20mM final)

1.0mL 0.1M Spermidine (10mM final)

600uL 0.5M PIPES, pH 7.4 (15.1g PIPES in 100mL goes into solution at pH 7.4)

5.0mL 100% Methanol (50% final)

MilliQ H_2_O up to 10mL

10% Formaldehyde

a) Microwave 18mL PBS in a 50mL conical (lid ajar) placed in a beaker with water until the water starts to boil
b) Add 2g PFA and 140uL 1M NaOH
c) Vortex carefully until the PFA dissolves
d) Add additional PBS to 20mL
e) Cool to 37C, then filter into 1mL aliquots and freeze

Tris-Triton Buffer (TTB)

1.0mL 1M Tris HCL, pH 7.4

100uL Triton-X-100

20uL 0.5M EDTA

MilliQ H_2_O up to 10mL

40x Borate Buffer

0.618g H_3_BO_3_

5.0mL 1M NaOH (required for H_3_BO_3_ to go into solution)

MilliQ H_2_O up to 10mL, pH 9.2

Heat to dissolve.

To make a 1x dilution, take 250uL 40x solution and bring final volume to 10mL.

20mM DTT

0.03g DTT

MilliQ H_2_O up to 10mL

Freeze in 600uL aliquots

Blocking Buffer

0.1g BSA (final concentration 1%)

25uL Triton-X-100

1200uL 2.5M glycine (final concentration 0.3M)

200uL 500mM imidazole (final concentration 10mM)

8.58mL PBS

Antibody-A Buffer

2.0mL 5x PBS

5.0mL 2% BSA

50uL Triton-X-100

250uL 2A% NaN_3_

20uL 0.5M EDTA

MilliQ H_2_O up to 10mL

Antibody-B Buffer

10.0mL Antibody-A Buffer

0.2% BSA
